# *BoskR* – Testing Adequacy of Diversification Models Using Tree Shape

**DOI:** 10.1101/2020.12.21.423829

**Authors:** Orlando Schwery, Brian C. O’Meara

## Abstract

The study of diversification largely relies on model-based approaches, estimating rates of speciation and extinction from phylogenetic trees. While a plethora of different models exist – all with different features, strengths and weaknesses – there is increasing concern about the reliability of the inference we gain from them. Apart from simply finding the model with the best fit for the data, we should find ways to assess a model’s suitability to describe the data in an absolute sense. The R package *BoskR* implements a simple way of judging a model’s adequacy for a given phylogeny using metrics for tree shape, assuming that a model is inadequate for a phylogeny if it produces trees that are consistently dissimilar in shape from the tree that should be analyzed. Tree shape is assessed via metrics derived from the tree’s modified graph Laplacian spectrum, as provided by *RPANDA*. We exemplify the use of the method using simulated and empirical example phylogenies. *BoskR* was mostly able to correctly distinguish trees simulated under clearly different models and revealed that not all models are adequate for the empirical example trees. We believe the metrics of tree shape to be an intuitive and relevant means of assessing diversification model adequacy. Furthermore, by implementing the approach in an openly available R package, we enable and encourage researchers to adopt adequacy testing into their workflow.

## Introduction

In the study of diversification, model-based estimation of diversification rates is probably the most common approach. Depending on the question, researchers will analyze diversification of their study group under different models and use some indicator for model fit (*e*.*g*. likelihood or Akaike Information Criterion (Akaike 1973)) to decide which model describes their data best and thus supports one or the other hypothesis. However, as those are only measures of relative fit, the best fitting model might still not actually be a good one to describe the data, hence the need for a way to assess its absolute fit. While we cannot assess a model’s actual absolute fit, since we are usually lacking knowledge of the absolute truth behind our empirical data, we can assess whether a candidate model is adequate for certain dataset, by exploring whether it can describe key properties of that data. This can be done by checking whether data generated under the candidate model is similar to the data to be tested with regards to said key properties.

Testing model adequacy this way has been attempted before for different aspects of building phylogenies: Sullivan and Swofford (1997) demonstrated that using an inadequate substitution model can lead to false conclusions about phylogenetic relationships; Huelsenbeck *et al*. (2001) and Bollback (2002) used Bayesian posterior predictive simulations to test the adequacy of substitution models, with the frequency of site patterns as the one test statistic compared, and Foster (2004) used it to specifically test the adequacy of substitution models with regards to compositional heterogeneity of the nucleotide frequencies across the tree; Brown and ElDabaje (2008) extended their adequacy test to include partitioning (vs. not), and Rodrigue *et al*. (2009) included site-dependence between codons; Brown (2014) took the approach a step further, generating trees from the posterior predictive sequences, relying on inference-based comparison instead of data-based comparison, and also demonstrating that his inferential test statistics outperform the multinomial one used by Bollback (2002); and finally Lewis *et al*. (2013) implement two new approaches that allow to dissect the overall verdict on substitution model adequacy.

Moving away from the substitution models, Duchêne *et al*. (2015) used posterior predictive simulations to test the adequacy of molecular clock models, also demonstrating the lower performance of a multinomial test statistic, and on a post-tree-inference state of phylogenetic analysis. Pennell *et al*. (2015) have introduced adequacy testing for models of continuous trait evolution and explored how it evaluates common trait evolution models across many empirical data sets.

With focus on diversification, Revell, Harmon and Glor (2005) have shown that using under-parametrized substitution models will affect the assessment of diversification dynamics (as inferred by the γ statistic of Pybus and Harvey (2000)), erroneously suggesting they are slowing down. Rabosky and Goldberg (2015) addressed the adequacy of the actual diversification models, in the specific case of models for trait dependent speciation, exploring aspects that render those models inadequate and cautioning against using them without additional testing, *e*.*g*. using neutral trait simulations. Another way of addressing the issue of model inadequacy was introduced with the inclusion of hidden states in state dependent diversification models (Beaulieu & O’Meara 2016; Caetano, O’Meara & Beaulieu 2018), which allow for the signal of rate heterogeneity to be identified in the tree through these hidden states, instead of erroneously assigning it to the observed traits through accidental imperfect association. Finally, following the example of earlier attempts to test the adequacy of substitution models, Höhna, May and Moore (2015) explicitly highlighted capabilities to use posterior-predictive simulations for adequacy testing in their R package *TESS* (suggesting to use γ). However, the habit of actually testing model adequacy has not yet found its place in the standard procedures of diversification studies (Brown & ElDabaje 2008).

Here, we provide a way of assessing a model’s adequacy using tree shape as a comparative metric. The dynamics of speciation and extinction determine the shape of a phylogeny, by affecting its topology and branch length distribution. In fact, these are the core data we use when estimating diversification rates. Different modes of diversification – described by different diversification models – should thus manifest in a range of different tree shapes, some of which are unique to a specific mode of diversification, whereas others are shared. Thus, if a model cannot generate phylogenies of similar shape to a given empirical phylogeny, it is probably not an adequate model to describe its underlying diversification process.

## Material and Methods

### Implementation of Adequacy Test

To assess adequacy, we rely on tree shape metrics derived from the tree’s graph Laplacian (Lewitus & Morlon 2016) as implemented in the package *RPANDA* (Morlon et al. 2016) in R (R Development Core Team 2014). The method constructs a modified graph Laplacian from a phylogeny and creates a spectral density profile from its eigenvalues. The main aspects of tree shape are characterized from three properties of the spectral density profile including: principal Eigenvalue (λ*; an indicator for species richness and phylogenetic diversity), asymmetry/skewness (ψ; indicating stemminess vs. tippiness), and peak height/kurtosis (η; indicating tree balance). A more detailed description of the method can be found in Lewitus and Morlon (2016).

The procedure to assess the adequacy of any diversification model for a given phylogeny then works as follows: First we calculate the shape metrics (λ*, ψ, η) of the empirical tree, as well as the rate estimates under the model being assessed. Next, we simulate a set of phylogenies under the same model using the same rates. We then assess the shape metrics for each of the resulting simulated trees, to gain a model- and parameter-specific distribution for each metric. The adequacy of a model for the empirical tree can then be assessed by comparing the shape metrics of the empirical tree to those of the simulated ones. This can be done in a number of ways, the following of which were tried: corrected or uncorrected p-values, 2D or 3D convex hulls, and Euclidean distances between and among simulated and empirical trees. A comparison of the results yielded by each method, suggested Bonferroni-corrected p-values and 2D convex hulls as a combination that provided a good compromise between performance and computational burden. In the first case, we calculate a p-value for each shape metric, based on the empirical distribution function of the metrics describing the simulated trees, and then apply a Bonferroni-correction to the three p-values. For the second method, we construct convex hulls around the points made up of a combination of two metrics describing the simulated trees. The resulting polygons from a minimal set of lines that connect the outermost points of the set and including all points of that set. We then test whether the point corresponding to the empirical tree lies within the convex hull or not, doing so for all three pairwise combinations of metrics. It is worth noting that if the cloud of simulated points has a non-convex shape, a potential could exist where the convex hull includes a significant amount of metric space that is not actually occupied by any of the simulated trees, but could be by the empirical tree.

The assumption is that a model which does not create trees of comparable shape to the empirical phylogeny is inadequate to describe that phylogeny: there is some aspect of the real data that is not predicted by the model, suggesting that a key biological process is not included. For an adequate model, one would expect the empirical tree to fall within the point cloud of the simulated trees, and thus to not significantly fall outside of the simulated distribution for each metric. If there is a significant difference between any of the shape properties of empirical and simulated trees, one has to assume that the model is not adequate, as in that there are processes underlying the generation of the empirical tree that are not accounted for by the employed model. While the focus here is on ultrametric trees of extant taxa, without the extinct lineages, it is worth noting that both the calculation of Laplacian spectra and the simulation of trees under a given model are generally possible, allowing the inclusion of extinct taxa (fossils) in the tree, if information on their placement is available.

The two methods of assessing whether the significant differences exist are expected to complement each other in two ways: the convex hull method tends to be more inclusive, only considering a model inadequate if the empirical tree metrics are entirely outside of the cloud of simulated tree metrics, whereas the Bonferroni-corrected p-values are more strict, potentially deeming a model inadequate if the empirical metrics lie within the simulated cloud of points, but not falling within 95% of those values, thus only sharing similar shape features with a few simulation-outliers. On the other hand, the p-values are univariate, thus while an empirical tree’s shape metrics could lie within 95% of each separate metric of the simulated trees, the convex hulls could reveal whether that particular combination of two metrics occurs within the simulations. Visually inspecting the results in a three-dimensional scatterplot can give us a more intuitive understanding of the data and can *e*.*g*. help clarifying cases where the two methods (Bonferroni and convex hulls) do not agree. Inspecting how exactly the tree shapes differ should allow one to explore where the inadequacy arises. An overview of all functions in the package can be found in Table 1.

**Table 1:** Functions available in *BoskR*.

| Function | Description |
| --- | --- |
| CombineTrees | combines several phylo objects into required format |
| GetMetricTreeSets | simulates sets of trees under a certain model using parameters estimated from an empirical tree |
| GetTreeMetrics | obtains shape metrics for sets of trees |
| GetTreeParams | estimates diversification model parameters of a set of trees |
| PvalMetrics | calculates a p-value for the empirical tree metrics based on the empirical cumulative density function of the simulated metrics |
| ScatterMetrics | generates a 3D-scatterplot of empirical and simulated tree metrics |
| TreeCorr | tests whether trees are ultrametric, bifurcating, and ordered cladewise, and corrects automatically if possible |
| plotPvalMetricsCDF | plots the empirical metrics and obtained p-values on their respective cumulative distribution functions |
| plotPvalMetricsPDF | plots empirical metrics on distribution of simulated metrics |

### Test of Specificity

To assess whether the method works reliably, we simulated 300 trees under a constant rate birth-death model, and subsequently tested whether said model is adequate to analyze those trees. Using *sim*.*bd*.*taxa* from the R package *TreeSim* (Stadler 2011), we simulated 300 trees with 200 extant taxa each, pruned extinct lineages, with diversification rates set to λ=1 and μ=0.2. As described above, we then for each tree estimated their rate parameters, simulated 1000 trees each with those parameters under a constant rate birth-death model, inferred the tree shape metrics for all trees and compared those of the sets of simulated 1000 trees with those of the initial trees they are based on.

### Examples of Use

#### Simulated Examples

First, we simulate ten trees each under a constant rate birth-death model and a birth-death model with mass extinctions and rate shifts. Both tree sets were simulated using the R package *TreeSim* (Stadler 2011), using *sim*.*bd*.*taxa* and *sim*.*rateshift*.*taxa* respectively. The goal was to obtain two sets of trees with shapes that are distinct enough to assure they do not result from the same processes, thus providing a positive control example. All trees were simulated to have 200 extant taxa, be completely sampled, and not include extinct lineages. The diversification rates for the pure birth-death simulations were set to λ=1 and μ=0.2. The other set of trees started with λ=1.5 and μ=0.1 up to a mass extinction event at time 4.5, which only 0.1 of all lineages survived, followed by λ=0.2 and μ=0.1. We then subjected the trees to the same procedure as above to test whether the constant rate birth-death model is adequate for them.

#### Empirical Examples

Further, we use a set of three published empirical phylogenies including a tree of 87 whale species (Steeman *et al*. 2009), a tree of 77 species of Ericaceae (derived from Schwery *et al*. 2015), and a tree of 11 species of *Calomys* (Pigot, Owens & Orme 2012), all of which are part of the example tree set implemented in the package. Using three of the currently implemented models – constant rate birth-death, time-dependent birth-death with exponential λ and constant μ, and diversity dependent birth-death with linear λ and constant μ – we inferred their rate estimates, simulated 1000 trees with those estimates under each model, inferred tree shape metrics for all trees and compared the empirical trees with their corresponding sets of simulated trees.

## Results

### Test of Specificity

Of the 300 trees simulated under a constant rate birth-death model, 287 ran successfully, that is, without any issues that prevented the rate estimation or tree simulation. Of the 13 trees that failed to run, 10 failed completely (suggesting issues at the stage of rate estimation), while three only failed to calculate the metrics of the simulated trees. Among the successful runs, the constant rate birth-death model was deemed adequate for 283 (98.6%) of which using Bonferroni corrected p-values (α=0.0139), and for 285 (99.3%) of which using 2D convex hulls (α=0.007). We expected the constant rate birth-death model, as the generating model, to be adequate for all trees, though of course there is a chance that a rare simulated tree will incorrectly reject the true generating model. Of the sets of trees for which birth-death was inadequate, only one of the two identified through 2D convex hulls was also among those identified through p-values, potentially demonstrating the value of using them complementary. It is also worth noting that when only run with 100 simulations per empirical tree, p-values inferred the model correctly as adequate at a similar rate (98.3%), while 2D convex hulls performed much worse (85.5%), perhaps due to few points leading to bad estimation of the shape of the point cloud (with just 100 points, their number at the outline is very small with large spaces between them).

### Examples of Use

#### Simulated Examples

As would be expected, most trees simulated under birth-death fell well within the range of shape metrics of their corresponding simulations, whereas the shape metrics of all ten trees simulated with mass extinction events fell significantly outside the range of their simulations for at least one metric (Table 2). However, the birth-death model also turned out to be inadequate for one of the initial birth-death trees (BD 2, see Table 2). Both methods of assessing adequacy (Bonferroni-corrected p-values and 2D convex hulls) agreed in their verdict of which trees the birth-death model was adequate for.

**Table 2:**
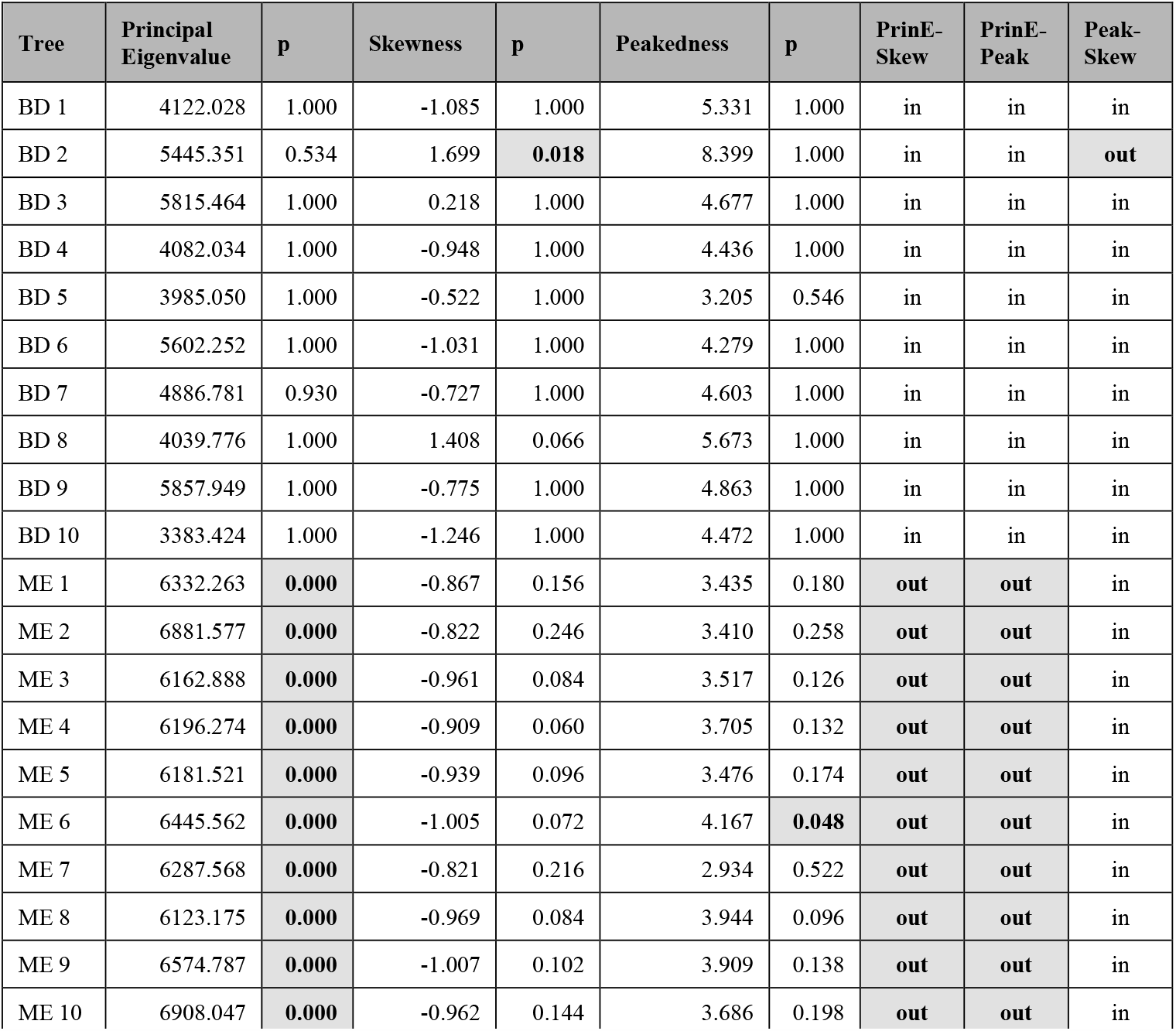
Results simulated examples. For each initially simulated tree, the three shape metrics (principal Eigenvalue, skewness, peakedness) are shown, followed by their corresponding Bonferroni-corrected p-value. The last three columns indicate for each pairwise combination of two metrics whether the initial tree is inside or outside of the polygon around the values of the simulated trees. P-values below 0.05 and points outside of 2D polygons are in bold and highlighted gray. BD= birth-death, ME= mass extinction, PrinE= principal Eigenvalue, Skew= skewness, Peak= peakedness.

| Tree | Principal Eigenvalue | p | Skewness | p | Peakedness | p | PrinE-Skew | PrinE-Peak | Peak-Skew |
| --- | --- | --- | --- | --- | --- | --- | --- | --- | --- |
| BD 1 | 4122.028 | 1.000 | -1.085 | 1.000 | 5.331 | 1.000 | in | in | in |
| BD 2 | 5445.351 | 0.534 | 1.699 | **0.018** | 8.399 | 1.000 | in | in | **out** |
| BD 3 | 5815.464 | 1.000 | 0.218 | 1.000 | 4.677 | 1.000 | in | in | in |
| BD 4 | 4082.034 | 1.000 | -0.948 | 1.000 | 4.436 | 1.000 | in | in | in |
| BD 5 | 3985.050 | 1.000 | -0.522 | 1.000 | 3.205 | 0.546 | in | in | in |
| BD 6 | 5602.252 | 1.000 | -1.031 | 1.000 | 4.279 | 1.000 | in | in | in |
| BD 7 | 4886.781 | 0.930 | -0.727 | 1.000 | 4.603 | 1.000 | in | in | in |
| BD 8 | 4039.776 | 1.000 | 1.408 | 0.066 | 5.673 | 1.000 | in | in | in |
| BD 9 | 5857.949 | 1.000 | -0.775 | 1.000 | 4.863 | 1.000 | in | in | in |
| BD 10 | 3383.424 | 1.000 | -1.246 | 1.000 | 4.472 | 1.000 | in | in | in |
| ME 1 | 6332.263 | **0.000** | -0.867 | 0.156 | 3.435 | 0.180 | **out** | **out** | in |
| ME 2 | 6881.577 | **0.000** | -0.822 | 0.246 | 3.410 | 0.258 | **out** | **out** | in |
| ME 3 | 6162.888 | **0.000** | -0.961 | 0.084 | 3.517 | 0.126 | **out** | **out** | in |
| ME 4 | 6196.274 | **0.000** | -0.909 | 0.060 | 3.705 | 0.132 | **out** | **out** | in |
| ME 5 | 6181.521 | **0.000** | -0.939 | 0.096 | 3.476 | 0.174 | **out** | **out** | in |
| ME 6 | 6445.562 | **0.000** | -1.005 | 0.072 | 4.167 | **0.048** | **out** | **out** | in |
| ME 7 | 6287.568 | **0.000** | -0.821 | 0.216 | 2.934 | 0.522 | **out** | **out** | in |
| ME 8 | 6123.175 | **0.000** | -0.969 | 0.084 | 3.944 | 0.096 | **out** | **out** | in |
| ME 9 | 6574.787 | **0.000** | -1.007 | 0.102 | 3.909 | 0.138 | **out** | **out** | in |
| ME 10 | 6908.047 | **0.000** | -0.962 | 0.144 | 3.686 | 0.198 | **out** | **out** | in |

#### Empirical Examples

Both the birth-death and the density-dependent model are adequate for all three phylogenies tested, while the time-dependent model seems to only be adequate for Tree 3 (*Calomys*), as the other two trees fall outside the simulated distributions for skewness and peakedness, and outside of all three polygons (Table 3).

**Table 3:** Results empirical examples. For each of the three models, the three shape metrics (principal Eigenvalue, skewness, peakedness) of the empirical trees are shown, followed by their corresponding Bonferroni-corrected p-values. The last three columns indicate for each pairwise combination of two metrics whether the initial tree is inside or outside of the polygon around the values of the simulated trees. P-values below 0.05 and points outside of 2D polygons are in bold and highlighted gray. Tree 1= whales, Tree 2= Ericaceae, Tree 3= *Calomys*, BD= birth-death, TD= time dependent birth-death, DD= diversity dependent birth-death ME= mass extinction, PrinE= principal Eigenvalue, Skew= skewness, Peak= peakedness.

| Model | Tree | Principal Eigenvalue | p | Skewness | p | Peakedness | p | PrinE-Skew | PrinE-Peak | Peak-Skew |
| --- | --- | --- | --- | --- | --- | --- | --- | --- | --- | --- |
| BD | 1 | 11340.708 | 1.000 | 0.592 | 1.000 | 2.065 | 0.936 | in | in | in |
|  | 2 | 33280.806 | 1.000 | 0.129 | 1.000 | 1.989 | 0.738 | in | in | in |
|  | 3 | 111.085 | 1.000 | 0.710 | 1.000 | 1.389 | 1.000 | in | in | in |
| TD | 1 | 11340.708 | 0.108 | 0.592 | **0.000** | 2.065 | **0.000** | **out** | **out** | **out** |
|  | 2 | 33280.806 | 0.150 | 0.129 | **0.000** | 1.989 | **0.000** | **out** | **out** | **out** |
|  | 3 | 111.085 | 1.000 | 0.710 | 1.000 | 1.389 | 1.000 | in | in | in |
| DD | 1 | 11340.708 | 1.000 | 0.592 | 0.786 | 2.065 | 0.462 | in | in | in |
|  | 2 | 33280.806 | 1.000 | 0.129 | 1.000 | 1.989 | 0.336 | in | in | in |
|  | 3 | 111.085 | 1.000 | 0.710 | 1.000 | 1.389 | 1.000 | in | in | in |

## Discussion

We implemented a way to test model adequacy of diversification models, by comparing tree shape metrics (derived from Laplacian spectra) of a tree with those of trees simulated under the same model and parameters. Models are not adequate to describe the diversification process underlying a tree, if those models lead to trees of significantly different shape. We tested the approach on sets of simulated and empirical trees to demonstrate its use.

An advantage of using *BoskR* for empirical data is it allows us seeing which aspects of shape cause the mismatch of a tree with a model. When inspecting the source of the inadequacy of the birth-death model for the mass extinction trees, it becomes apparent that the principal Eigenvalue is the source of mismatch. (Table 2). The actual values of metrics (Table 4) show that the high principal Eigenvalue of the mass extinction tree tends to be greater than that of the birth-death trees, and that the trees simulated from their parameters under the birth-death model have consistently and markedly lower principal Eigenvalues. The principal Eigenvalue of phylogenetic Laplacian spectra is an indicator for species richness and phylogenetic diversity (Lewitus & Morlon 2016). And indeed, while all initial trees had 200 tips, the simulated trees derived from the birth-death set had an average of 357.95 tips (min: 4, max: 3798), whereas those derived from the mass extinction set had an average of 20.46 tips (min: 2, max: 157).

**Table 4:** Metrics of Simulated Trees. For all three shape metrics (Principal Eigenvalue, Skewness, Peakedness) the values for each initially simulated tree are shown followed by the mean, minimal and maximal values of their corresponding set of 1000 trees simulated with the same parameters. BD= birth-death, ME= mass extinction.

| Tree | Principal Eigenvalue |  |  |  | Skewness |  |  |  | Peakedness |  |  |  |
| --- | --- | --- | --- | --- | --- | --- | --- | --- | --- | --- | --- | --- |
|  | initial | mean | min | max | initial | mean | min | max | initial | mean | min | max |
| BD 1 | 4122.028 | 3544.249 | 66.293 | 16677.736 | -1.085 | -0.471 | -2.134 | 3.858 | 5.331 | 4.743 | 1.124 | 11.297 |
| BD 2 | 5445.351 | 21648.058 | 522.668 | 103476.154 | 1.699 | -1.051 | -2.504 | 2.338 | 8.399 | 7.706 | 1.589 | 17.410 |
| BD 3 | 5815.464 | 6948.965 | 239.777 | 31975.499 | 0.218 | -0.553 | -2.648 | 3.337 | 4.677 | 5.515 | 1.354 | 15.076 |
| BD 4 | 4082.034 | 4400.988 | 108.527 | 20366.263 | -0.948 | -0.611 | -2.173 | 2.782 | 4.436 | 5.119 | 1.501 | 13.664 |
| BD 5 | 3985.050 | 7054.063 | 169.758 | 31808.846 | -0.522 | -0.820 | -2.293 | 2.928 | 3.205 | 5.927 | 1.477 | 15.543 |
| BD 6 | 5602.252 | 9924.124 | 228.569 | 49798.910 | -1.031 | -0.795 | -2.470 | 3.539 | 4.279 | 6.166 | 1.250 | 16.263 |
| BD 7 | 4886.781 | 13639.045 | 326.075 | 77492.732 | -0.727 | -0.984 | -2.426 | 2.193 | 4.603 | 7.057 | 1.690 | 17.792 |
| BD 8 | 4039.776 | 8845.887 | 106.573 | 51451.470 | 1.408 | -0.883 | -2.214 | 3.219 | 5.673 | 6.296 | 1.160 | 15.784 |
| BD 9 | 5857.949 | 9063.837 | 182.765 | 43736.546 | -0.775 | -0.665 | -2.471 | 3.940 | 4.863 | 5.924 | 1.167 | 16.875 |
| BD 10 | 3383.424 | 2497.903 | 74.447 | 13926.700 | -1.246 | -0.471 | -2.034 | 2.973 | 4.472 | 4.329 | 1.042 | 12.513 |
| ME 1 | 6332.263 | 590.048 | 45.513 | 2921.191 | -0.867 | 0.457 | -1.176 | 2.889 | 3.435 | 1.872 | 0.756 | 4.579 |
| ME 2 | 6881.577 | 796.181 | 49.063 | 5629.196 | -0.822 | 0.339 | -1.431 | 3.124 | 3.410 | 2.033 | 0.735 | 7.614 |
| ME 3 | 6162.888 | 550.735 | 44.277 | 3147.839 | -0.961 | 0.472 | -1.096 | 3.131 | 3.517 | 1.873 | 0.745 | 6.045 |
| ME 4 | 6196.274 | 538.247 | 44.712 | 2565.612 | -0.909 | 0.479 | -1.277 | 2.930 | 3.705 | 1.875 | 0.728 | 25.277 |
| ME 5 | 6181.521 | 599.297 | 43.749 | 2577.140 | -0.939 | 0.432 | -1.309 | 2.773 | 3.476 | 1.918 | 0.789 | 7.058 |
| ME 6 | 6445.562 | 698.300 | 46.356 | 3618.554 | -1.005 | 0.368 | -1.345 | 2.635 | 4.167 | 1.995 | 0.719 | 4.997 |
| ME 7 | 6287.568 | 620.810 | 44.693 | 2482.020 | -0.821 | 0.387 | -1.268 | 3.198 | 2.934 | 1.935 | 0.720 | 5.692 |
| ME 8 | 6123.175 | 658.443 | 43.344 | 2809.674 | -0.969 | 0.377 | -1.203 | 3.419 | 3.944 | 2.020 | 0.731 | 6.413 |
| ME 9 | 6574.787 | 709.712 | 45.960 | 4468.763 | -1.007 | 0.360 | -1.300 | 2.687 | 3.909 | 2.039 | 0.721 | 11.556 |
| ME 10 | 6908.047 | 752.806 | 49.307 | 3701.842 | -0.962 | 0.432 | -1.177 | 3.719 | 3.686 | 2.012 | 0.827 | 8.317 |

While counting the number of tips might have been sufficient in this particular case, it is worth noting that a difference in principal Eigenvalue between two trees could also be found if they had the same number of taxa. For the birth-death set, the estimated diversification rates were λ=1.014 and μ=0.254, but only λ=0.233 and μ=0 for the mass extinction set, leading to the difference in number of tips. The reason behind these differences derives from a constant-rate birth-death model generating (or assuming) an even distribution of speciation and extinction events per lineage over time. The continuous lineage accumulation in the initial birth-death tree set could be described adequately that way, but the mass extinction trees had a high lineage accumulation initially that plateaued after the mass extinction and the subsequent lower net diversification. This inflated the birth-death rate estimates, which cannot adequately represent this kind of dynamic.

The single birth-death tree for which the birth-death model turned out inadequate (BD2), differed significantly from the simulated set of 1000 trees in skewness (Table 2). It shows a higher (positive) skewness (1.699) than all other initial trees (Table 4), while the associated simulated trees have similar asymmetries as the simulation sets based on the other initial birth-death trees (−2.504 to 2.338, mean of −1.051). Skewness describes the distribution of branching events over time, with negative values implying stemmy trees, and positive values implying tippy trees (Lewitus & Morlon 2016). Thus, BD2 is supposedly tippier than the other initial trees and those simulated from its parameters. When inspecting the tree and its associated Laplacian spectrum (Figure 1), this tree stands out by having a small peak of higher Eigenvalues next to the main peak, which shifts the skewness of the whole spectrum to be higher, and which should correspond to the long branch clade at the root of the tree (sister to all the other taxa). A clade like this – with not just the diversity difference, but also the long-branch connection to the other taxa – would be rather uncommon under a constant rate birth-death tree but can occur by chance. Thus, it would be equally rare to get a similar-looking tree within the simulated set of 1000 birth-death trees, leading to the result that the model is inadequate, despite the initial tree being simulated under a birth-death model. Indeed, when dropping this clade from the tree and rerunning the adequacy test, birth-death comes out as adequate for this tree.

**Figure 1:**
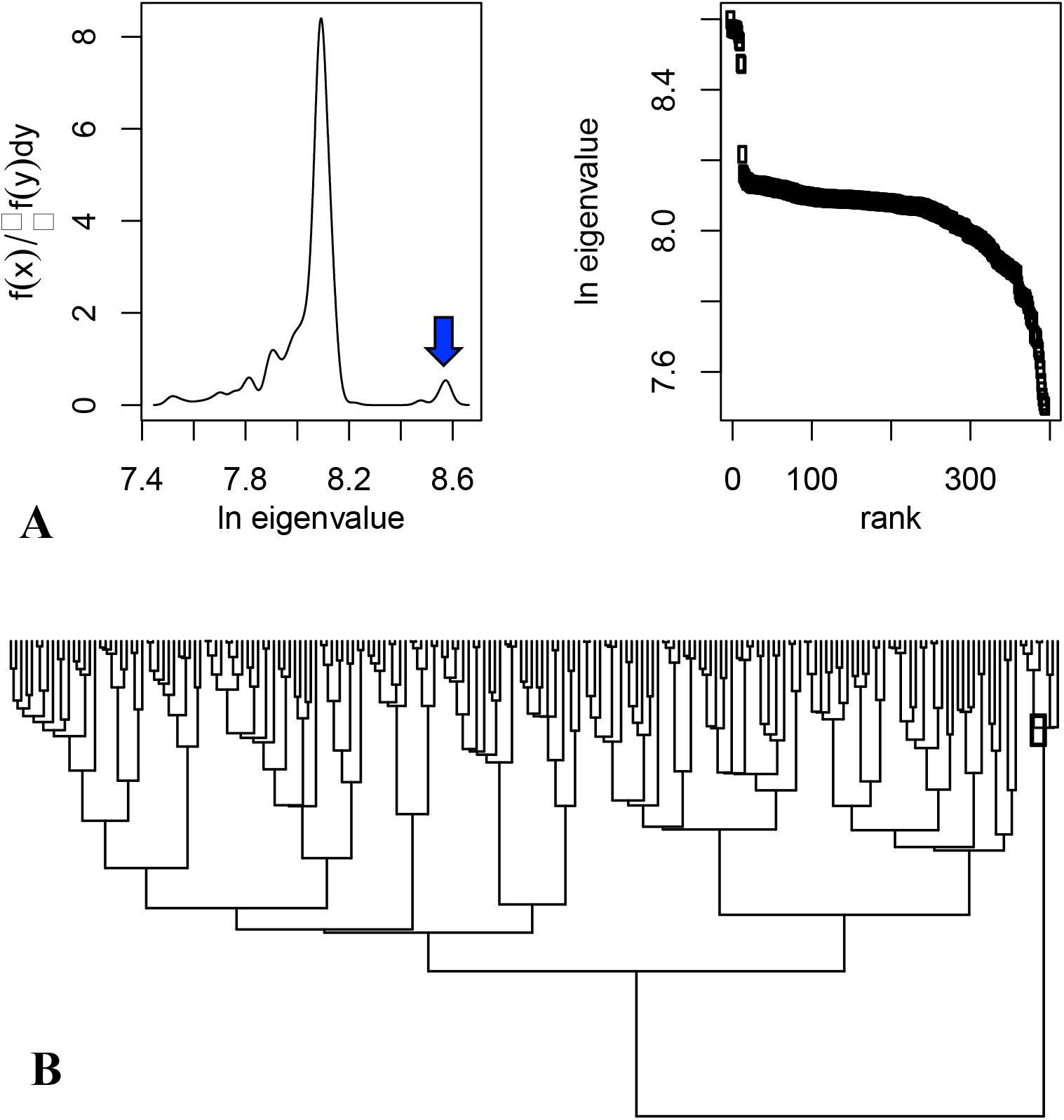
Phylogeny BD2 and associated Laplacian spectrum. **A:** Laplacian spectrum of the tree BD2, left, and eigenvalues sorted by size, right. **B:** The phylogeny BD2. The blue dot and arrow indicate the clade (and its associated peak in the spectrum) that caused the shift in skewness.

For the empirical examples, a comparison of Trees 1 and 2 and the tree sets derived from them under the time-dependent model, reveals that they mainly disagree in skewness and peakedness (Table 5). As mentioned above, skewness describes whether a tree is tippy or stemmy, while peakedness describes how evenly the Eigenvalues are distributed, with low values indicating a homogeneous distribution, thus balanced trees, and high values accordingly imbalanced trees (Lewitus & Morlon 2016). For the two trees where the time dependent model is inadequate, both skewness and peakedness are higher for the empirical trees than for the 1000 simulated ones derived from them (Table 5). Thus, they should be more tippy and imbalanced than they would be under a time dependent process. Indeed, when looking at the actual trees, it becomes apparent that the simulated trees are extraordinarily stemmy and balanced when compared to their empirical counterpart (

**Table 5:** Metrics of Empirical Trees. For all three shape metrics (principal Eigenvalue, skewness, peakedness), the values for each empirical tree are shown followed by the mean, minimal and maximal values of their corresponding set of 1000 trees simulated with the same parameters under all three models. BD= birth-death, TD= time dependent birth-death, DD= diversity dependent birth-death.

| Model | Tree | Principal Eigenvalue |  |  |  | Skewness |  |  |  | Peakedness |  |  |  |
| --- | --- | --- | --- | --- | --- | --- | --- | --- | --- | --- | --- | --- | --- |
|  |  | initial | mean | min | max | initial | mean | min | max | initial | mean | min | max |
| BD | 1 | 11340.708 | 10491.411 | 320.914 | 47380.279 | 0.592 | -0.218 | -1.733 | 3.467 | 2.065 | 3.447 | 0.781 | 7.907 |
|  | 2 | 33280.806 | 33454.247 | 981.070 | 181480.025 | 0.129 | -0.228 | -1.733 | 3.779 | 1.989 | 3.418 | 0.952 | 8.982 |
|  | 3 | 111.085 | 115.525 | 14.961 | 565.026 | 0.710 | 0.809 | -1.207 | 3.432 | 1.389 | 1.716 | 0.723 | 13.604 |
| TD | 1 | 11340.708 | 3851.769 | 179.289 | 17380.147 | 0.592 | -0.090 | -0.235 | 0.498 | 2.065 | 1.190 | 0.756 | 1.728 |
|  | 2 | 33280.806 | 12583.687 | 586.423 | 78178.138 | 0.129 | -0.052 | -0.311 | 0.000 | 1.989 | 1.013 | 0.735 | 1.510 |
|  | 3 | 111.085 | 106.331 | 14.961 | 435.420 | 0.710 | 0.377 | -0.756 | 2.497 | 1.389 | 1.413 | 0.723 | 5.380 |
| DD | 1 | 11340.708 | 11447.491 | 554.804 | 24436.742 | 0.592 | -0.370 | -1.610 | 3.625 | 2.065 | 3.511 | 1.094 | 7.034 |
|  | 2 | 33280.806 | 33913.463 | 1483.834 | 65119.609 | 0.129 | -0.366 | -1.502 | 3.299 | 1.989 | 3.373 | 1.059 | 8.771 |
|  | 3 | 111.085 | 100.894 | 26.156 | 141.440 | 0.710 | 0.784 | -0.601 | 3.164 | 1.389 | 1.706 | 0.810 | 13.650 |

Figure 2). The diversification rates estimated from the empirical trees (and used as input for the simulated sets) show that extinction is close to zero for all three trees, and that the speciation rate of whales and Ericaceae decrease over time (Figure 3), which would be consistent with stemmy trees.

**Figure 2:**
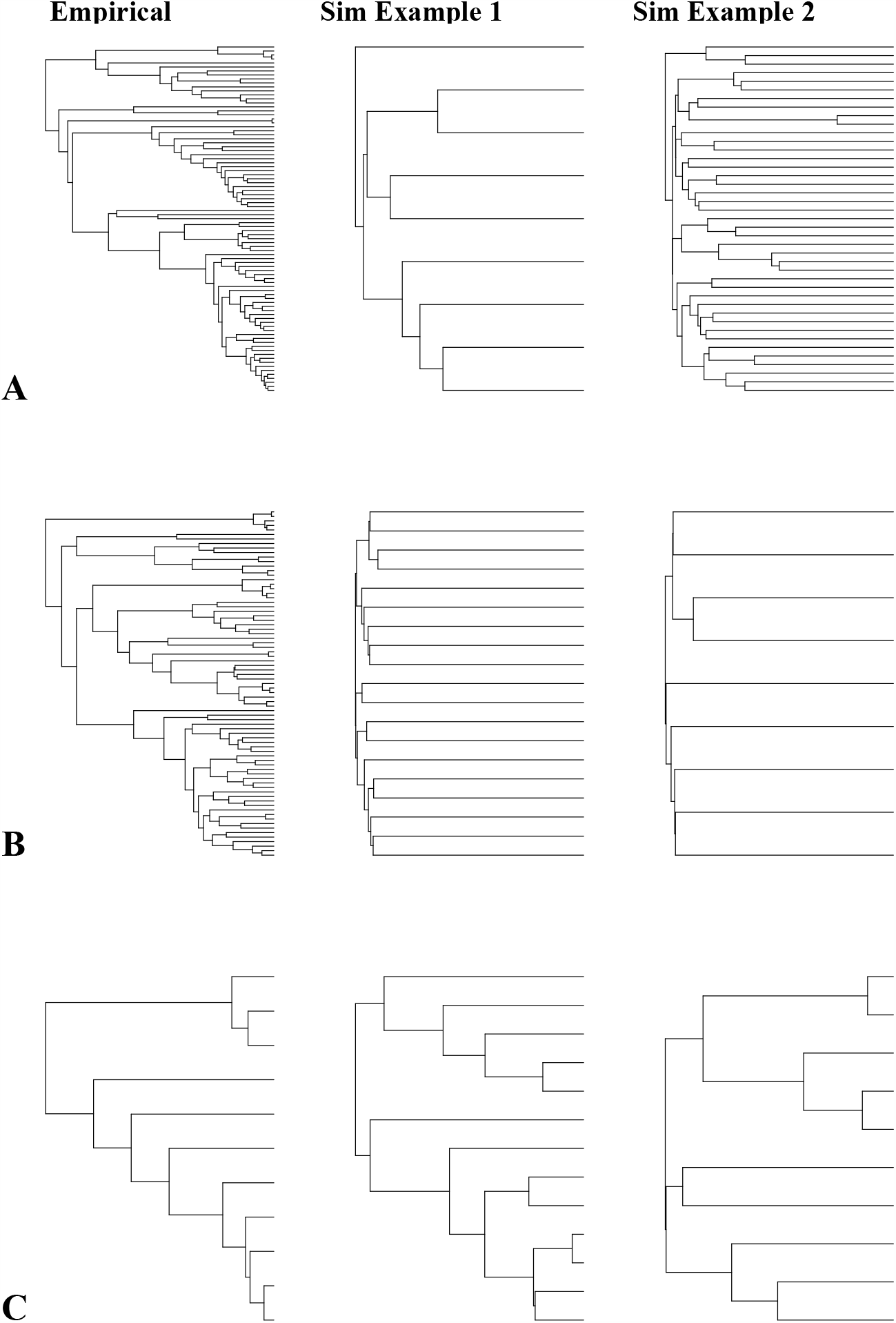
Empirical phylogenies and examples of associated simulations. For each empirical phylogeny, two example trees are shown, simulated with the same parameters as the empirical tree, and under the time dependent model. **A**: whales, **B**: Ericaceae, **C**: *Calomys*.

**Figure 3:**
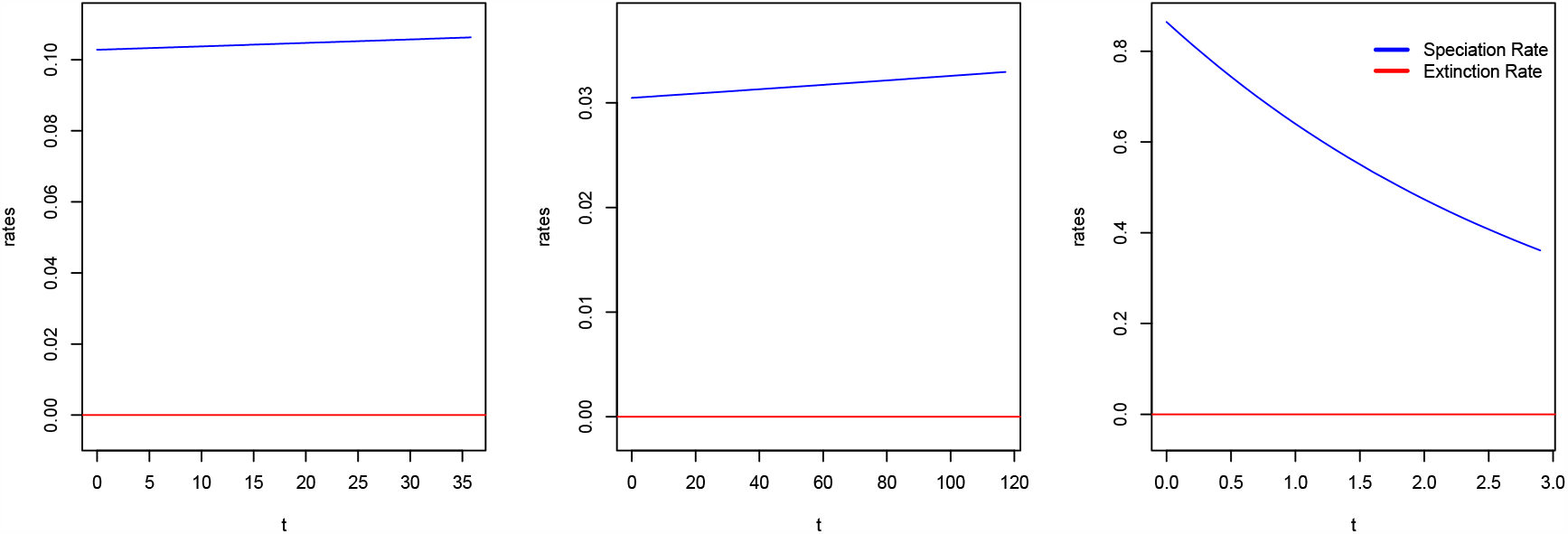
Diversification rates for empirical trees under time-dependent model. Speciation and extinction rates estimated from the empirical phylogenies under the time dependent model. **A**: whales, **B**: Ericaceae, **C**: *Calomys*, Blue: Speciation rate, red: Extinction rate.

We believe that *BoskR* will be a useful tool for researchers attempting to identify which models are adequate or inadequate to describe the diversification dynamics of a study group. These insights should allow for a more informed analysis and increased confidence in the obtained results. Since tree shape (being topology and branch lengths) is essentially the primary data we use in model-based inference of diversification rates, it should be an appropriate and intuitively relatable basis of assessing diversification model adequacy. Various other – arguably equally sensible – metrics of tree shape are of course available, and some of which might be assessed more easily. Indeed, for our simulated example, one could have simply used the number of tips to conclude that the birth-death model is inadequate for trees that evolved under a mass extinction model. However, number of tips would not have flagged BD2, which differed in skewness instead. Additionally, we have conditioned all simulations on crown age, which we deemed the most reasonable choice for most diversification questions. However, for cases where e.g. conditioning on number of surviving species, or both crown age and number of surviving species is more sensible, number of tips will not be a useful metric anymore. Using the Laplacian spectrum-based metrics in *BoskR* thus has the advantage to include several aspects of tree shape, and thereby also allows to explore whether and how one or several of these aspects render a model inadequate for a tree. Finally, by implementing the approach in an openly available R package, we enable and encourage researchers to adopt adequacy testing into their workflow.

It is, however, worth noting an implicit assumption of our approach: that the empirical tree under scrutiny is the true tree (or a reasonably close approximation of it). Our approach does not account for any errors and biases introduced in the tree building process. It would thus be possible that *i*.*e*. a group in fact diversified in a time-dependent fashion, while *BoskR* would deem this an inadequate model, because the signature of time dependence was lost during tree building and divergence time estimation. Furthermore, this approach does currently not account for issues in rate estimation. If the diversification rates are not estimated accurately to begin with, *BoskR* would likely label the corresponding model inadequate, without that necessarily being the case.

Future versions of the package will aim to include a wider range of models, to better reflect the diversity of models currently used in the field. However, as long as any given model is able to both estimate parameters and simulate trees under those parameters, the resulting outputs can be used as input for *BoskR* and can thus be analyzed without problem.

## Acknowledgements

We would like to express our thankfulness for helpful discussions and comments at various stages of this project to Athmanathan Senthilnathan, Lucas Santana-Souza, Jordan Bush, Luna Sanchez-Reyes, Jeremy Beaulieu, Matt Pennell, Dean Anderson, Emma Goldberg, Dan Simberloff, and Colin Sumrall.

## References

Akaike, H. (1973) Information theory and an extension of the maximum likelihood principle,[w:] proceedings of the 2nd international symposium on information, bn petrow, f. Czaki, Akademiai Kiado, Budapest.

Beaulieu, J.M. & O’Meara, B.C. (2016) Detecting Hidden Diversification Shifts in Models of Trait-Dependent Speciation and Extinction. Systematic Biology, 65, 583–601.

Bollback, J.P. (2002) Bayesian model adequacy and choice in phylogenetics. Molecular Biology and Evolution, 19, 1171–1180.

Brown, J.M. (2014) Detection of implausible phylogenetic inferences using posterior predictive assessment of model fit. Systematic Biology, 63, 334–348.

Brown, J.M. & ElDabaje, R. (2008) PuMA: Bayesian analysis of p artitioned (and u npartitioned) m odel a dequacy. Bioinformatics, 25, 537–538.

Caetano, D.S., O’Meara, B.C. & Beaulieu, J.M. (2018) Hidden state models improve state-dependent diversification approaches, including biogeographical models. Evolution, 72, 2308–2324.

Duchêne, D.A., Duchêne, S., Holmes, E.C. & Ho, S.Y. (2015) Evaluating the adequacy of molecular clock models using posterior predictive simulations. Molecular Biology and Evolution, 32, 2986–2995.

Foster, P.G. (2004) Modeling compositional heterogeneity. Systematic Biology, 53, 485–495.

Höhna, S., May, M.R. & Moore, B.R. (2015) TESS: an R package for efficiently simulating phylogenetic trees and performing Bayesian inference of lineage diversification rates. Bioinformatics, 32, 789–791.

Huelsenbeck, J.P., Ronquist, F., Nielsen, R. & Bollback, J.P. (2001) Bayesian inference of phylogeny and its impact on evolutionary biology. Science, 294, 2310–2314.

Lewis, P.O., Xie, W., Chen, M.-H., Fan, Y. & Kuo, L. (2013) Posterior predictive Bayesian phylogenetic model selection. Systematic Biology, 63, 309–321.

Lewitus, E. & Morlon, H. (2016) Characterizing and Comparing Phylogenies from their Laplacian Spectrum. Systematic Biology, 65, 495–507.

Morlon, H., Lewitus, E., Condamine, F.L., Manceau, M., Clavel, J. & Drury, J. (2016) RPANDA: an R package for macroevolutionary analyses on phylogenetic trees. Methods in Ecology and Evolution, 7, 589–597.

Pennell, M.W., FitzJohn, R.G., Cornwell, W.K. & Harmon, L.J. (2015) Model Adequacy and the Macroevolution of Angiosperm Functional Traits. American Naturalist, 186, E33–E50.

Pigot, A.L., Owens, I.P.F. & Orme, C.D.L. (2012) Speciation and Extinction Drive the Appearance of Directional Range Size Evolution in Phylogenies and the Fossil Record. Plos Biology, 10.

Pybus, O.G. & Harvey, P.H. (2000) Testing macro–evolutionary models using incomplete molecular phylogenies. Proceedings of the Royal Society of London. Series B: Biological Sciences, 267, 2267–2272.

R Development Core Team (2014) R: A language and environment for statistical computing. R Foundation for Statistical Computing, Vienna, Austria.

Rabosky, D.L. & Goldberg, E.E. (2015) Model Inadequacy and Mistaken Inferences of Trait-Dependent Speciation. Systematic Biology, 64, 340–355.

Revell, L.J., Harmon, L.J. & Glor, R.E. (2005) Under-parameterized model of sequence evolution leads to bias in the estimation of diversification rates from molecular phylogenies. Systematic Biology, 54, 973–983.

Rodrigue, N., Kleinman, C.L., Philippe, H. & Lartillot, N. (2009) Computational methods for evaluating phylogenetic models of coding sequence evolution with dependence between codons. Molecular Biology and Evolution, 26, 1663–1676.

Schwery, O., Onstein, R.E., Bouchenak-Khelladi, Y., Xing, Y., Carter, R.J. & Linder, H.P. (2015) As old as the mountains: the radiations of the Ericaceae. New Phytologist, 207, 355–367.

Stadler, T. (2011) Simulating Trees with a Fixed Number of Extant Species. Systematic Biology, 60, 676–684.

Steeman, M.E., Hebsgaard, M.B., Fordyce, R.E., Ho, S.Y.W., Rabosky, D.L., Nielsen, R., Rahbek, C., Glenner, H., Sorensen, M.V. & Willerslev, E. (2009) Radiation of Extant Cetaceans Driven by Restructuring of the Oceans. Systematic Biology, 58, 573–585.

Sullivan, J. & Swofford, D.L. (1997) Are guinea pigs rodents? The importance of adequate models in molecular phylogenetics. Journal of Mammalian Evolution, 4, 77–86.

